# Increased *S. epidermidis* in the airway-gut microbiome of infants with bronchopulmonary dysplasia

**DOI:** 10.64898/2026.04.03.715941

**Authors:** Zenna Solomon, Megan Eno, Sharon C. Thompson, Stephanie L. Rager, Jenny C. Jin, Melody Y. Zeng, Divya Keerthy, Stefan Worgall, Elizabeth L. Johnson, Andrea Heras

**Affiliations:** Department of Pediatrics, Weill Cornell Medicine, New York, NY; Division of Nutritional Sciences, Cornell University, Ithaca, NY; Howard Hughes Medical Institute, Cornell University, Ithaca, NY; Drukier Institute for Children’s Health, Weill Cornell Medicine, New York, NY; Neonatal and Perinatal Medicine, NewYork-Presbyterian Queens, Flushing, NY; Department of Genetic Medicine, Weill Cornell Medicine, New York, NY

## Abstract

**Rationale:** Bronchopulmonary dysplasia (BPD), the lung disease associated with premature birth, is a significant health problem, often with long-term respiratory consequences. Recent research has highlighted the potential role of the lung and gut microbiome in the development and progression of BPD, yet it is unclear what aspects of the microbiome may contribute to BPD susceptibility.

**Objectives:** To comprehensively characterize the lung and gut microbiomes of preterm infants and identify shared microbial taxa that are associated with BPD development.

**Methods:** Tracheal aspirate and stool samples were collected from 39 premature infants over the first month of life. To assess the taxonomic microbial composition of the lung and gut, samples were analyzed using shotgun metagenomic sequencing. BPD classification was determined using the National Institute of Child Health and Human Development severity-based definition at 36 weeks postmenstrual age.

**Measurements and Main Results:** Microbial communities of the lung and gut were significantly different between infants who went on to develop BPD and those who did not, with an enrichment of skin-associated microbial genera such as *Staphylococcus, Corynebacterium,* and *Cutibacterium* in infants who developed BPD. Specifically, *Staphylococcus epidermidis* was enriched in premature infants who developed BPD and was the most prominent species shared between lung and gut communities. Temporal changes in gut microbial communities co-occurred with feeding practices and antibiotic exposure, suggesting an influence of external factors on microbiome composition.

**Conclusions:** Our findings provide evidence that certain microbial colonization patterns among premature infants are closely associated with the pathogenesis and progression of BPD.

## Introduction

Bronchopulmonary dysplasia (BPD) is among the most prevalent complications of prematurity, and its incidence continues to increase as advances in neonatal care improve survival among the most vulnerable preterm infants.^1^ Infants who develop BPD may experience lifelong respiratory complications such as impaired lung function, airway hyperreactivity, and early development of chronic obstructive pulmonary disease.^2–4^ BPD is characterized by impaired alveolar and vascular development, influenced by antenatal and postnatal factors, including placental dysfunction, ventilator induced lung injury, and infections.^5^ However, the precise mechanisms by which these factors contribute to BPD pathogenesis remain incompletely understood.

Alterations in the development of the gut and lung microbiomes may contribute to abnormal physiological responses and play a key role in BPD pathogenesis.^6,7^ The gut and lungs interact bidirectionally (gut-lung axis), with the microbiome and its metabolites influencing the health of both organs.^7,8^ The lung and gut of an infant harbor a unique and dynamic microbiome that can be influenced by external factors.^9,10^ The early life of a preterm infant is a “critical window” during which time, health-associated microbiomes may influence the developing immune system. Deviations from the normal development of these health-associated microbial communities are commonly defined as a state of dysbiosis and can have implications for respiratory health.^6^ Specifically, dysbiosis may diminish lung epithelial barrier function, increase susceptibility to infections, and promote inflammation, all of which could result in lung tissue damage and contribute to BPD development and progression.^7,8^

Preterm birth results in early life exposure to many clinical factors, such as antibiotics, total parenteral nutrition, and mechanical ventilation, that alter typical microbiome development.^6,11^ These variable and dynamic clinical factors have hindered efforts to define the “normal” microbiome of a preterm infant creating challenges to the identification of specific microbiome alterations that contribute to BPD development. In general, preterm infants have a microbiome that is less diverse when compared to full-term infants.^12^ Evidence regarding the lung or gut microbiota of preterm infants is varied, with few cohorts profiled prior to BPD diagnosis. Moreover, reliance on high-level taxonomic summaries limits species-level inference and has primarily implicated broad taxa (e.g., *Ureaplasma,*^13^ *Actinetobacter,*^14^ *Staphylococcus,*^15^ *Proteobacteria*^10^). Furthermore, no studies have evaluated concurrent changes in gut and lung microbiome composition that contribute to BPD development at the resolution that allows for tracing of species sharing between the lung and gut environment. This study aims to comprehensively characterize the lung and gut microbiomes of preterm infants, identify shared microbial taxa that contribute to the development of BPD, and evaluate key microbiome modifiers, including feeding practices and antibiotic exposure.

## Materials and Methods

### Study design and subject enrollment

Preterm infants (born less than 37 weeks of gestation) were recruited from the level IV neonatal intensive care unit (NICU) of NewYork-Presbyterian/Weill Cornell Medicine in New York City, New York, between 2022 and 2024. Patients were categorized into 3 groups based on BPD status and intubation history: BPD and intubated (BPD^+^ Int^+^), no BPD and intubated (BPD^-^ Int^+^), and no BPD and no intubation (BPD^-^ Int^-^). BPD status was determined using the National Institute of Child Health and Human Development (NICHD) definition at 36 weeks postmenstrual age. Clinical data, including demographic information, antibiotic exposure, and feeding practices, were collected from the electronic health record.

### Tracheal aspirate and stool collection

Tracheal aspirates (TA) were collected daily during the first week of life and subsequently weekly until 30 days of life or until extubation, whichever occurred first. Stool samples were scheduled for daily collection during the first week of life and then weekly until 30 days of life. Variability in stool patterns in premature infants necessitated collection when samples were available therefore not every infant had daily/weekly samples. Stool samples were collected from 36 infants and TA samples were collected from 21 infants of which samples from 10 infants contained microbial reads. For all samples, day of life at the time of collection was defined such that the date of birth corresponded to day of life 0, with day of life 1 beginning 24 hours after birth, consistent with standard NICU nomenclature. See **Supplementary Methods** for additional details.

### DNA extraction and metagenomic library preparation and sequencing

DNA extraction from stool and TA samples was performed using the Qiagen DNeasy PowerSoil Pro Kit (Qiagen, #47016) following the manufacturer’s instructions. Libraries were prepared using the NexteraXT DNA Library Preparation Kit (Illumina). See **Supplementary Methods** for details.

### Species and clinical isolate tracking analysis

Stool and TA samples were assessed for bacterial species sharing between the metagenomes of these body sites using Sourmash (v4.9.3). Stool and TA metagenomes were aligned to reference genomes (listed in Table S1) using minimap2 (v2.30-r1287). For clinical isolate tracking analysis, putative *Staphylococcus epidermidis* strains were sequenced using Oxford Nanopore long read sequencing (Plasmidasaurus) and bacterial genomes were mapped to stool and TA metagenomes for profiling. See **Supplementary Methods** for details.

### Statistical analysis

All statistical analysis was performed using RStudio version 2024.04.1+748 and GraphPad Prism version 10.2.3. Microbiome processing and visualization were performed using the R packages phyloseq, tidyverse, ggplot2, vegan, dplyr, and magrittr.^16,17^ Differences in relative abundance between infant groups were assessed using a Kruskal-Wallis test. For species level analyses, Benjamini-Hochberg (BH) false discovery rate (FDR) procedure was used for multiple hypothesis testing. Pairwise-Wilcoxon tests were used for post-hoc analysis of significant results. Post-hoc comparisons were considered significant when Benjamini-Hochberg FDR adjusted p-values were < 0.05. The following significance codes were used for plotting: **** < 0.0001, *** < 0.001, ** < 0.01, and * < 0.05.

### Study approval

The study was approved by the Weill Cornell Medicine Institutional Review Board (IRB protocol #22-05024765 and #20-04021833).

## Results

### Study demographics

Race, ethnicity, and gender distributions were similar between groups (Table 1). Infants with BPD who were intubated (BPD^+^ Int^+^) were more likely to be extreme preterm and to have extremely low birthweight compared to infants who were intubated without BPD (BPD^-^ Int^+^), and infants without BPD (BPD^-^ Int^-^) (**Table 1**).

**Table 1.** Demographics.

| Characteristic | BPD- Int- | BPD- Int+ | BPD+ Int+ | p-value<sup>2</sup> | BPD- Int- vs. BPD- Int+ | BPD- Int- vs. BPD+ Int+ | BPD- Int+ vs. BPD+ Int+ |
| --- | --- | --- | --- | --- | --- | --- | --- |
|  | N = 12<sup>1</sup> | N = 10<sup>1</sup> | N = 17<sup>1</sup> |  |  |  |  |
| **Gender** |  |  |  | 0.5 | >0.9 | 0.8 | >0.9 |
| Female | 7 (58%) | 5 (50%) | 6 (35%) |  |  |  |  |
| Male | 5 (42%) | 5 (50%) | 11 (65%) |  |  |  |  |
| **Gestational Age (weeks)** | 34.15 (33.65, 34.25) | 31.45 (30.20, 33.10) | 27.00 (25.20, 29.40) | <0.001 | 0.01 | <0.001 | <0.001 |
| **Gestational Age Group** |  |  |  | <0.001 | 0.005 | <0.001 | 0.005 |
| Extreme Preterm <28 Weeks | 0 (0%) | 0 (0%) | 10 (59%) |  |  |  |  |
| Very Preterm 28-32 Weeks | 1 (8.3%) | 7 (70%) | 7 (41%) |  |  |  |  |
| Moderate Preterm 32-34 Weeks | 2 (17%) | 2 (20%) | 0 (0%) |  |  |  |  |
| Late Preterm 34-37 Weeks | 9 (75%) | 1 (10%) | 0 (0%) |  |  |  |  |
| **Multigestational Pregnancy** | 4 (33%) | 2 (20%) | 2 (12%) | 0.3 | >0.9 | 0.6 | >0.9 |
| **Birth Weight (g)** | 2,080 (1,895, 2,290) | 1,565 (1,310, 1,950) | 870 (690, 1,170) | <0.001 | 0.01 | <0.001 | <0.001 |
| **Birth Weight Group** |  |  |  | <0.001 | 0.056 | <0.001 | 0.001 |
| Extremely Low (<1000g) | 0 (0%) | 0 (0%) | 11 (65%) |  |  |  |  |
| Very Low (1000g - 1500g) | 1 (8.3%) | 5 (50%) | 5 (29%) |  |  |  |  |
| Low (1500g - 2500g) | 11 (92%) | 5 (50%) | 1 (5.9%) |  |  |  |  |
| **Ethnicity** |  |  |  | 0.054 | 0.8 | 0.8 | 0.078 |
| Hispanic | 2 (20%) | 0 (0%) | 7 (41%) |  |  |  |  |
| Not Hispanic | 8 (80%) | 10 (100%) | 10 (59%) |  |  |  |  |
| Unknown | 2 | 0 | 0 |  |  |  |  |
| **Race** |  |  |  | 0.6 | >0.9 | >0.9 | >0.9 |
| Asian | 3 (30%) | 5 (50%) | 3 (19%) |  |  |  |  |
| Black | 3 (30%) | 2 (20%) | 7 (44%) |  |  |  |  |
| Mixed/Other | 3 (30%) | 1 (10%) | 2 (13%) |  |  |  |  |
| White | 1 (10%) | 2 (20%) | 4 (25%) |  |  |  |  |
| Unknown | 2 | 0 | 1 |  |  |  |  |
| **Mode of Delivery** |  |  |  | 0.9 | >0.9 | >0.9 | >0.9 |
| Cesarean Section | 9 (75%) | 8 (80%) | 14 (82%) |  |  |  |  |
| Vaginal | 3 (25%) | 2 (20%) | 3 (18%) |  |  |  |  |
| **Chorioamnionitis** | 0 (0%) | 1 (10%) | 2 (12%) | 0.6 | >0.9 | >0.9 | >0.9 |
| **Maternal Preeclampsia** | 1 (8.3%) | 2 (20%) | 8 (47%) | 0.066 | 0.6 | 0.13 | 0.5 |

<sup>1</sup>n (%); Median (Q1, Q3)
<sup>2</sup>Fisher's exact test; Kruskal-Wallis rank sum test

### *Staphylococcus epidermidis* is the most prevalent species in the lower airway microbiome of infants who develop BPD

Shotgun metagenomic sequencing of tracheal aspirates (TA) revealed low microbial biomass. Of the 84 samples sequenced across 21 infants, microbial reads were detected in 29 samples from 10 infants, all with BPD (Figure 1). Across samples with detected microbial reads, 18 unique bacterial species were identified (Figure 2C). The most prevalent bacterial species, *Staphylococcus epidermidis,* was present in five out of ten infants with TA sequencing reads (Figure 2C). Other microbial species identified in two or more infants included *Bacteroides thetaiotaomicron, Escherichia coli, Klebsiella pneumoniae, Staphylococcus lugdunensis*, and *Streptococcus mitis* (Figure 2C). The infant lung microbiome was typically dominated by one to two microbial species. Longitudinal analysis of the seven (out of 10) infants with multiple TA samples demonstrated temporal shifts in microbial composition (Figure 2A). The relative abundance of each taxon was averaged across all TA samples at the genus level (Figure 2B). Across all samples, *Staphylococcus* had the highest average relative abundance across the cohort, followed by *Klebsiella, Streptococcus, Escherichia*, and *Bacteroides* (Figure 2B). Overall, detectable lung microbial communities were limited to infants who developed BPD and were characterized by low diversity and dominance by a few taxa.

**Figure 1.**
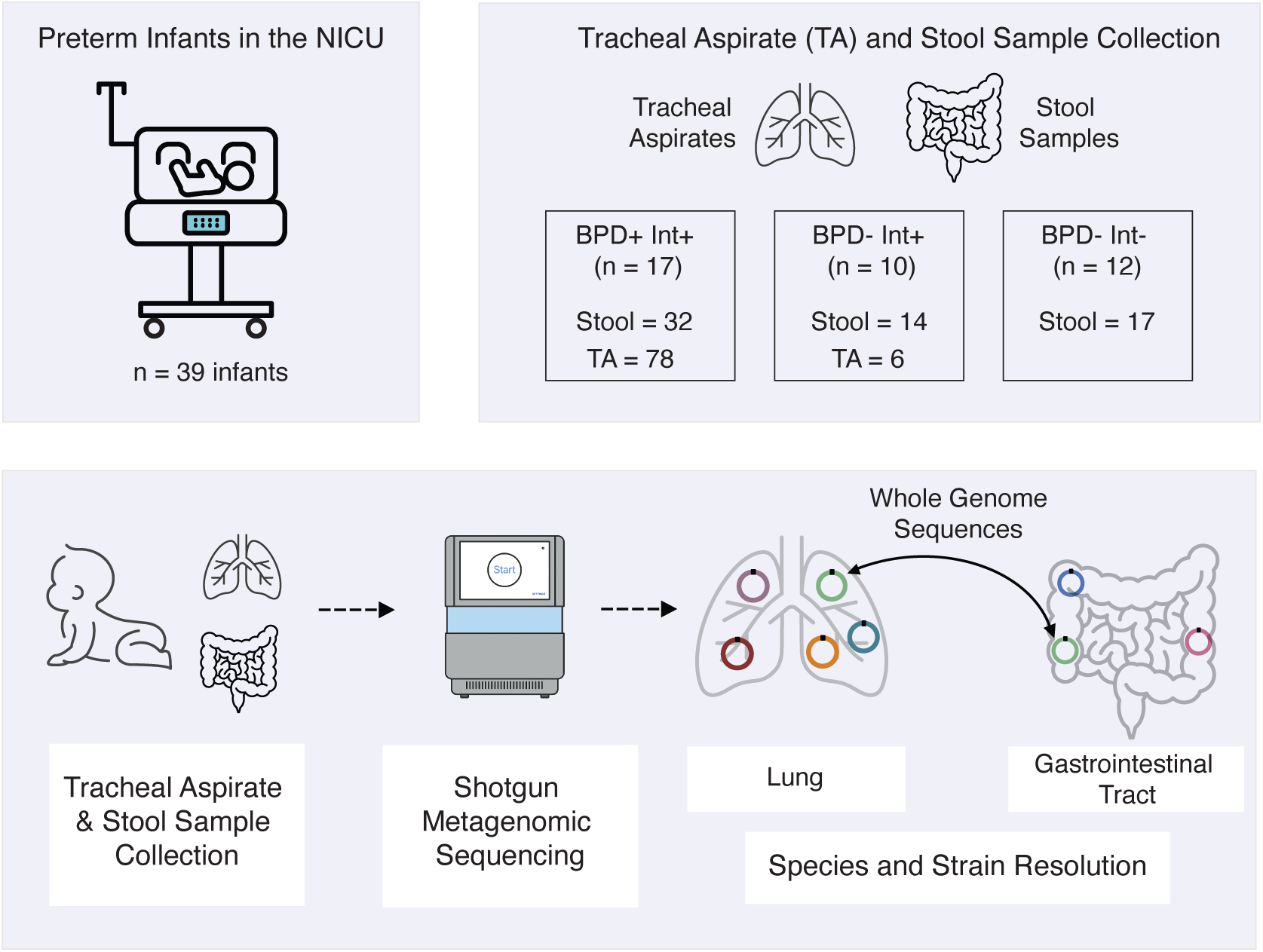
Schematic of study design and experimental procedures. Diagrams showing total number of infants in the study, number of infants in each disease/procedure state, number of samples collected of each sample type and the experimental procedures used to analyze samples. Neonatal intensive care unit (NICU), intubated (Int), bronchopulmonary dysplasia (BPD).

**Figure 2.**
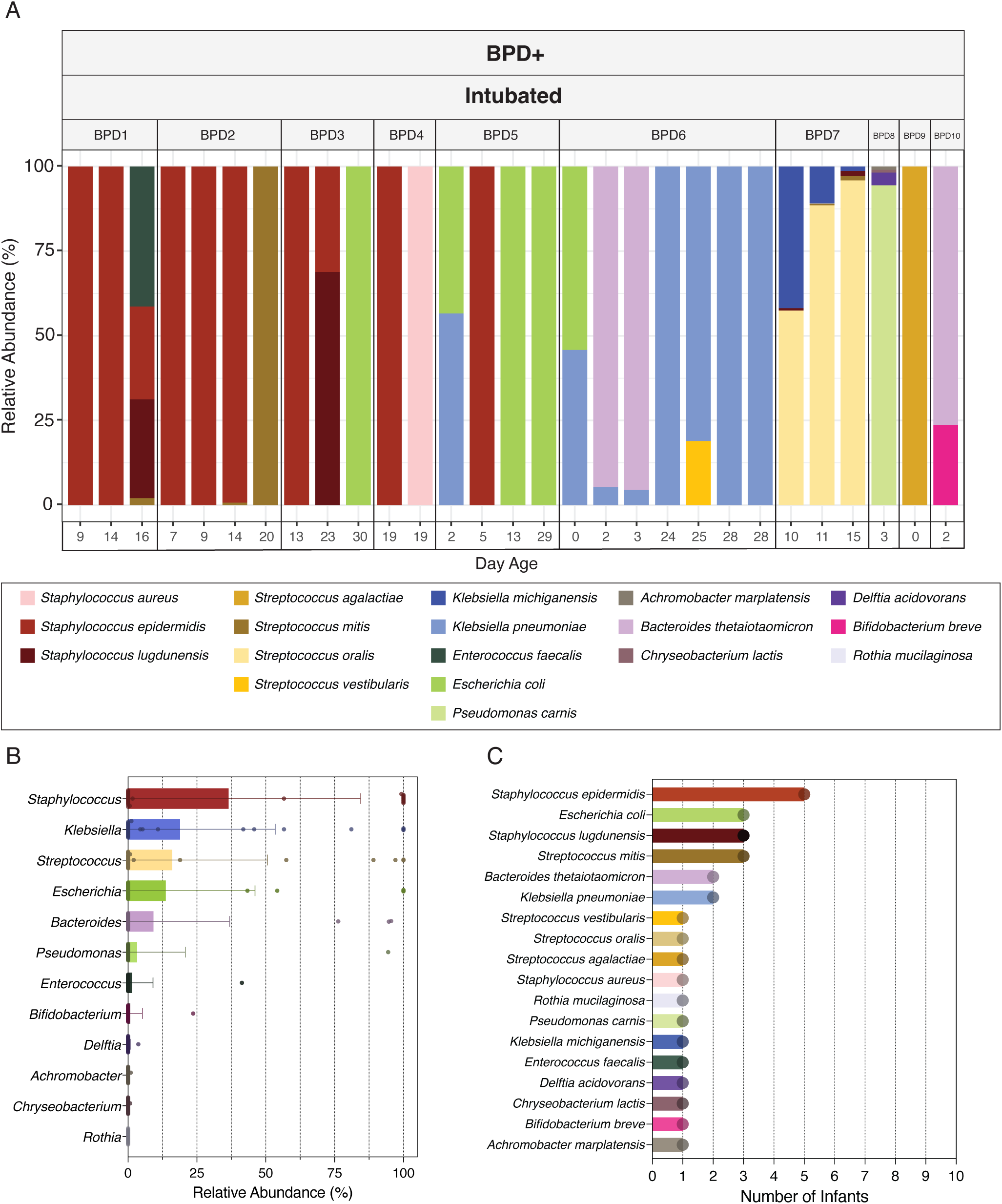
Microbial composition of the lung microbiome in infants that developed BPD. (A) Relative abundance of microbial species in tracheal aspirate samples determined by shotgun metagenomic sequencing. Plots are separated by infant. (B) Bar plots displaying mean relative abundance of microbial genera across all tracheal aspirates. Error bars = standard deviation [n=29] (C) Occurrence of microbial species in the lung microbiome of preterm, intubated infants, based on tracheal aspirate samples from 10 infants with microbial reads.

### Gut microbiome composition is influenced by BPD and intubation status

Shotgun metagenomic sequencing was used to profile the gut microbiome composition of 36 infants (Figure 3A). Infants were stratified into three groups, with one to five stool samples collected per infant: (1) BPD^+^ Int^+^, n = 32, (2) BPD^-^ Int^+^, n = 14, and (3) BPD^-^ Int^-^, n = 17. The total abundance of known skin-associated genera,^18^ which include *Staphylococcus, Corynebacterium*, and *Cutibacterium* was compared between groups (Figure 4A). A Kruskal-Wallis rank-sum test demonstrated a significant difference between groups (p = 0.005). Post-hoc analysis using BH adjusted pairwise Wilcoxon rank sum tests demonstrated a significant difference in the microbial abundance of skin-associated microbes between BPD^+^ Int^+^ infants compared to both BPD^-^ Int^+^ (p-adj = 0.048) and BPD^-^ Int^-^ infants (p-adj = 0.008). There was no significant difference between BPD^-^ Int^+^ and BPD^-^ Int^-^ infants (p-adj = 0.706).

**Figure 3.**
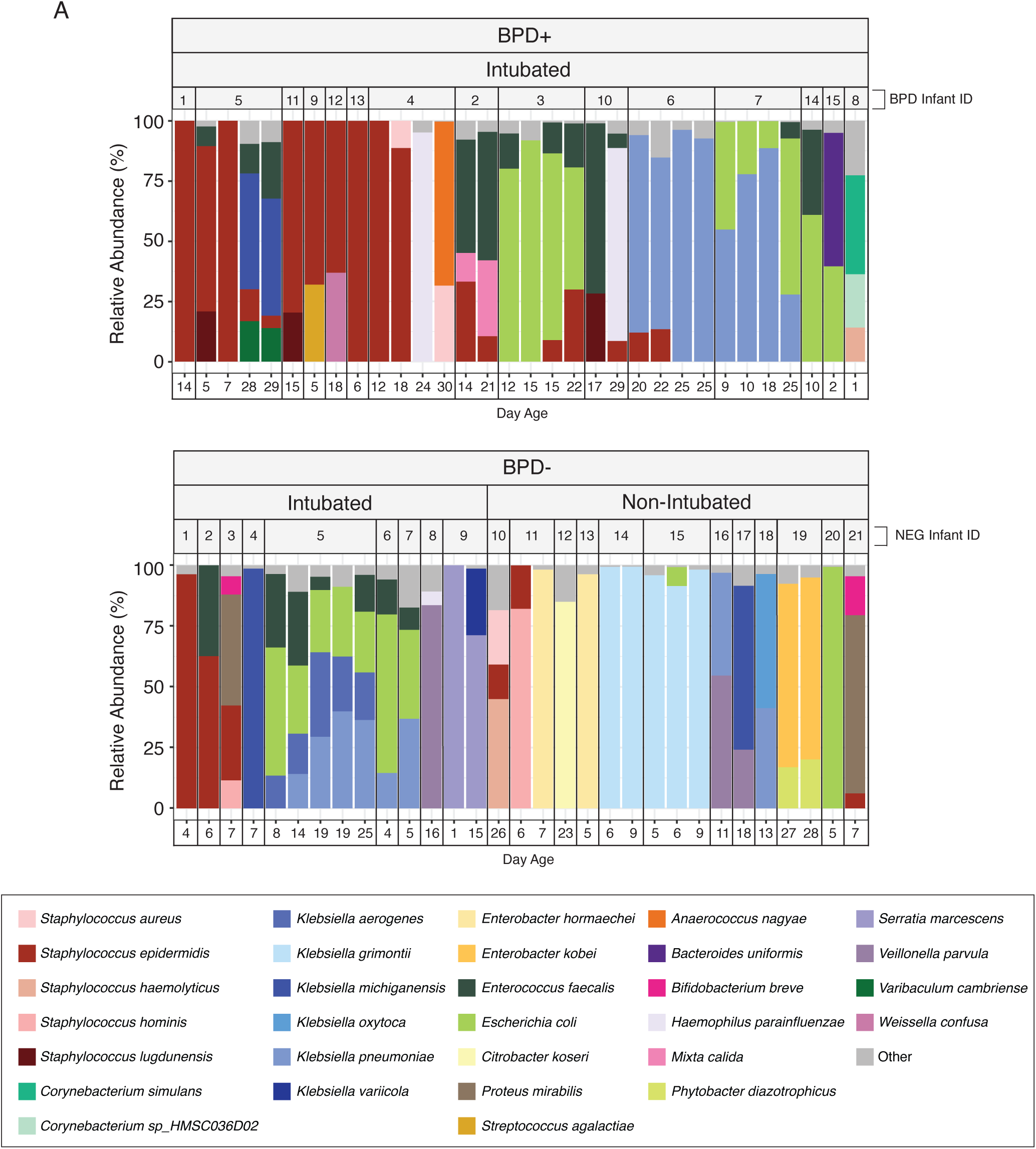
Gut microbiome composition of preterm infants across BPD and intubation statuses. Comparison of the relative abundance of microbial taxa across 3 groups: 1. BPD-Positive Intubated (BPD^+^ Int^+^), 2. BPD-Negative Intubated (BPD^-^ Int^+^), and 3. BPD-Negative Non-Intubated (BPD^-^ Int^-^). For visualization, species accounting for less than 5% relative abundance in any given sample and species with <20% relative abundance across all samples were grouped into the ‘Other’ category.

**Figure 4.**
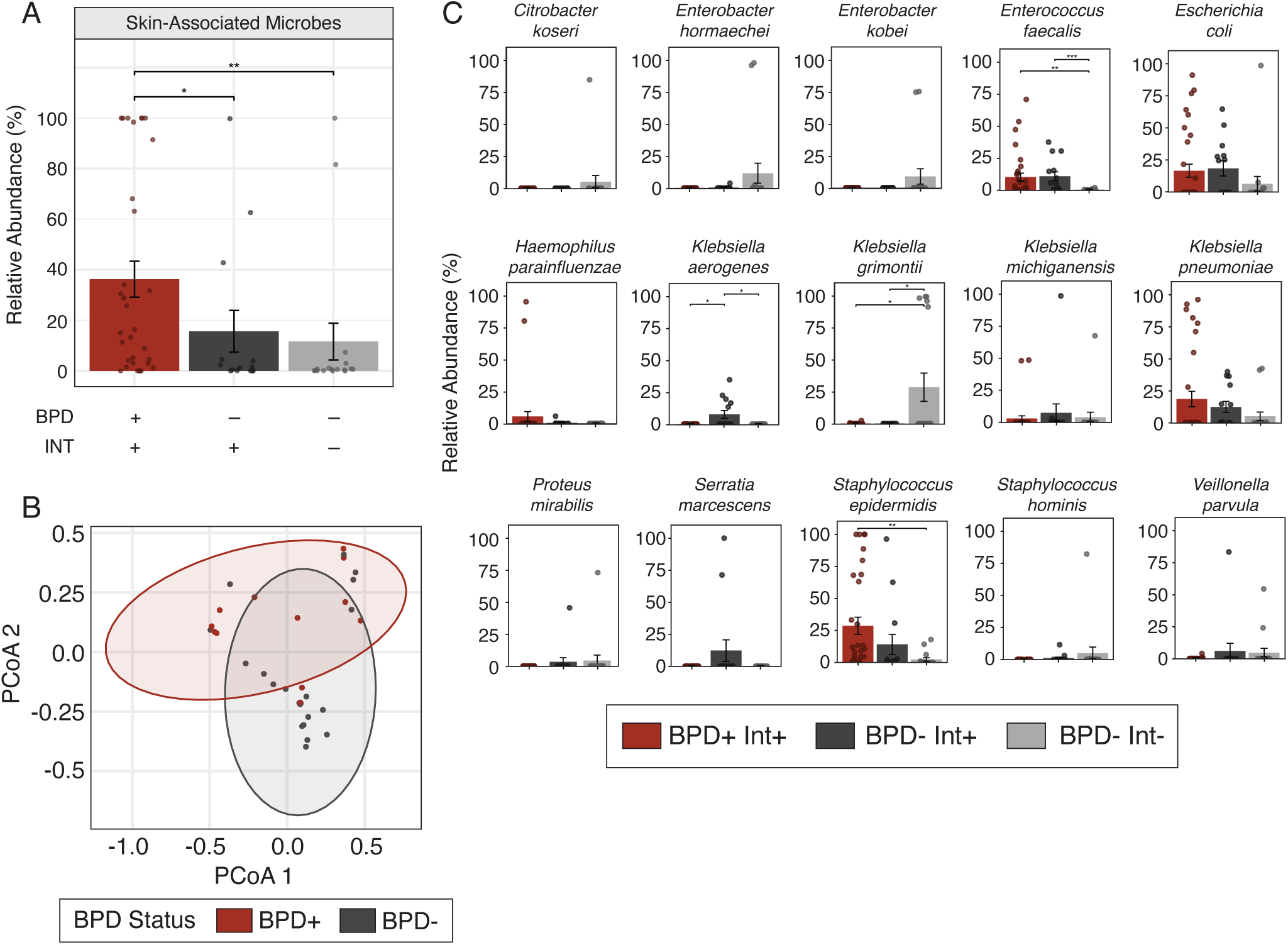
Differences in microbial communities and individual taxa across BPD and intubation statuses. (A) Average abundance of skin-associated genera including *Corynebacterium*, *Cutibacterium*, and *Staphylococcus*, compared between groups (Kruskal-Wallis, p = 0.005) [n: BPD^+^ Int^+^ = 32 samples across 15 infants, BPD^-^ Int^+^ = 14 samples across 9 infants, BPD^-^ Int^-^ = 17 samples across 12 infants].(B) Beta diversity determination using Bray-Curtis dissimilarity between BPD^+^ and BPD^-^ infants (PERMANOVA: R^2^ = 0.061, F = 2.226, p = 0.021) [n: BPD^+^ Int^+^ = 15, BPD^-^ Int^+^ = 9, BPD^-^ Int^-^ = 12].(C) Comparison of the fifteen most abundant species in the microbiome between groups of differing BPD and intubation status using a Kruskal-Wallis tests with BH FDR procedure and Wilcoxon pairwise tests for post-hoc analysis [n: BPD^+^ Int^+^ = 32 samples across 15 infants, BPD^-^ Int^+^ = 14 samples across 9 infants, BPD^-^ Int^-^ = 17 samples across 12 infants].

To assess potential differences in infant gut microbiome community structure between groups, principal coordinate analysis (PCoA) was performed using Bray-Curtis dissimilarity. A PERMANOVA test demonstrated microbial community composition significantly differed between BPD^+^ Int^+^, BPD^-^ Int^+^, and BPD^-^ Int^-^ infants (R^2^ = 0.094, F = 1.72, p = 0.022) (Supplementary Figure 1A). Comparison of BPD^+^ and BPD^-^ infants using a PERMANOVA revealed a significant difference between groups (R^2^ = 0.061, F = 2.226, p = 0.021) (Figure 4B). Comparison of Int^+^ and Int^-^ infants using a PERMANOVA also revealed a significant difference between groups (R^2^ = 0.072, F = 2.636, p = 0.007) (Supplementary Figure 1B). Regarding within sample alpha diversity metrics, Shannon diversity was compared among BPD^+^ Int^+^, BPD^-^ Int^+^, and BPD^-^ Int^-^ infants and a Kruskal-Wallis test revealed no significant differences between groups (χ^2^ = 0.625, df = 2, p = 0.73) (Supplemental Figure 1C). Comparison of Shannon diversity using a Wilcoxon rank sum test with continuity correction demonstrated no significant differences between BPD^+^ and BPD^-^ infants (W = 148, p = 0.773) (Supplemental Figure 1D) or Int^+^ and Int^-^ infants (W = 156, p = 0.6995) (Supplemental Figure 1E).

To better understand specific taxa driving differences in microbiome composition between conditions, the fifteen most abundant microbial species were identified and compared across the three infant groups (Figure 4C). Four species showed significant differences between groups: *Enterococcus faecalis* (p-adj = 0.011)*, Staphylococcus epidermidis* (p-adj = 0.020)*, Klebsiella aerogenes* (p-adj = 0.024), and *Klebsiella grimontii* (p-adj = 0.045). Post-hoc analysis for pairwise comparisons used Wilcoxon rank sum tests with BH adjustment. *Enterococcus faecalis* was lower in BPD^-^ Int^-^ infants compared to both BPD^+^ Int^+^ (p-adj = 0.0048) and BPD^-^ Int^+^ infants (p-adj = 0.0004). *Klebsiella aerogenes* was higher in BPD^-^ Int^+^ infants compared to both BPD^+^ Int^+^ (p-adj = 0.044) and BPD^-^ Int^-^ infants (p-adj = 0.011). *Klebsiella grimontii* was higher in BPD^-^ Int^-^ infants compared to both BPD^+^ Int^+^ (p-adj = 0.049) and BPD^-^ Int^+^ infants (p-adj = 0.049). Finally, *Staphylococcus epidermidis* was higher in BPD^+^ Int^+^ infants compared to BPD^-^ Int^-^ infants (p-adj = 0.004) Taken together, these results demonstrate significant differences in the abundance of key bacterial taxa and in overall community structure in infants that develop BPD, compared to that in preterm infants that do not develop BPD, indicating that BPD status may be linked to specific taxa of the gut microbiome in infants.

### Species sharing occurred between lung and stool metagenomes of BPD^+^ Int^+^ infants

To investigate the possibility of shared bacterial taxa between the sites of the lower airway and the infant gut, species sharing analysis was performed on the human depleted metagenomes of the TA and stool samples of BPD^+^ Int^+^ infants. From this kmer-based analysis, species sharing of *Staphylococcus epidermidis* was identified between the lung and gut microbiomes of five infants across multiple sampling days (Figure 5 A,D), while *Escherichia coli* was identified in two infants across individual sample sets (Figure 5A,E), and *Klebsiella pneumoniae* and *Streptococcus agalactiae* were identified in one infant each over multiple samples and a single sample set, respectively (Figure 5 F,G). Of the total of ten BPD^+^ Int^+^ infants, this analysis revealed 5/5 infants with *Staphylococcus epidermidis* in the lung share this species in the gut, 2/3 infants with *Escherichia coli* in the lung share this species in the gut, 1/1 with *Streptococcus agalactiae* share this species in the gut, and 1/2 infants with *Klebsiella pneumoniae* share this species in the gut (Figure 5H). To further characterize the relatedness of these shared species across TA and stool metagenomes, reference-based alignments of TA and stool metagenomes to a curated genomic database was utilized (Table S1). This analysis resulted in significant alignments of *Staphylococcus epidermidis* in three infants (Supplemental Figure 2A-C) and *Klebsiella pneumoniae* and *Streptococcus agalactiae* in one infant each (Supplemental Figure 2 D,E). From these alignments, SNP-based comparisons revealed greater genetic heterogeneity in shared *Klebsiella pneumoniae* strains (∼30,000 SNPs) compared to *Staphylococcus epidermidis* (∼10,000 SNPs), consistent with closer relatedness of the latter across lung-gut pairs (Supplemental Figure 2 A-E). Using these two approaches, we identified within-infant species sharing between lung and gut metagenomes in seven BPD^+^ Int^+^ infants (Figure 5). Across BPD^+^ Int^+^ infants, species-sharing analysis revealed overlapping lung-gut colonization, most commonly with *Staphylococcus epidermidis* (identified in five infants) with close relatedness, with fewer shared events involving *Klebsiella pneumoniae*, *Streptococcus agalactiae*, and *Escherichia coli*, indicating that select taxa can inhabit both sites within the same infant.

**Figure 5.**
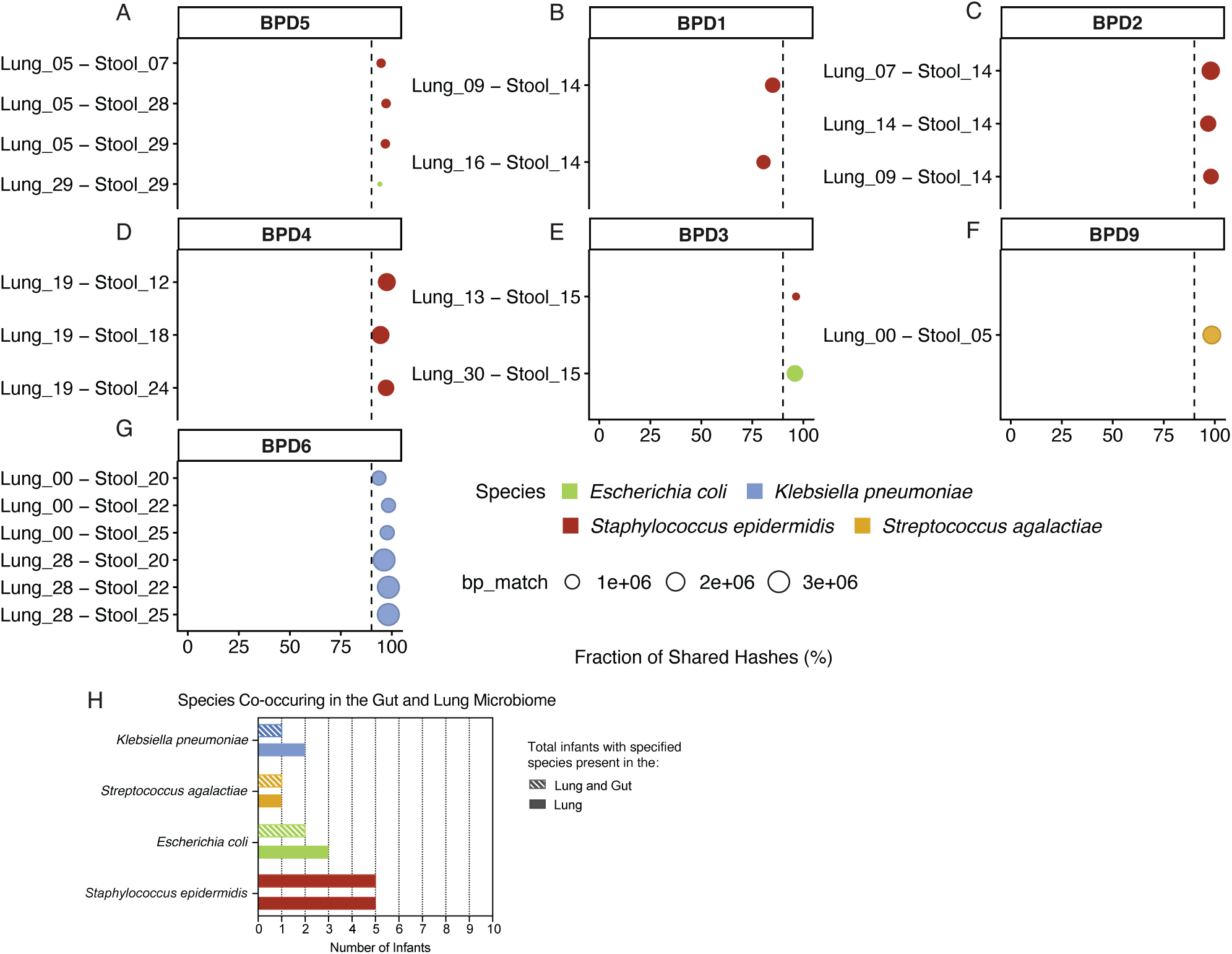
Species sharing analysis using Sourmash between lung and stool metagenomes in infants. (A) Infant – BPD5, (B) Infant – BPD1, (C) Infant – BPD2, (D) Infant – BPD4, (E) Infant – BPD3, (F) Infant – BPD9, and (G) Infant – BPD6 at different days of life. On the x-axis is the fraction of shared hashes between the lung and stool metagenomes and the given species. Also represented on the x-axis is a dotted line denoting a threshold of 90% shared hashes. Color of the point represents the species identified as shared between the lung and stool metagenomes, and size of the point represents the length of read aligned between lung, stool, and the given species genome. (H) Instances of microbial species sharing in the gut and lung microbiome of infants based on shared hashes.

### *Staphylococcus epidermidis* strain sharing occurred in BPD^+^ and BPD^-^ infants

*Staphylococcus epidermidis* AU12-03 is a clinical strain previously isolated from an intravascular catheter and previously identified in the NICU environment.^19^ *Staphylococcus epidermidis* AU12-03 (c70U7-1) was assessed for its presence in the lung and gut metagenomes where it was detected in seven stool samples from six infants but was not detected in any lung metagenomes (Figure 6A). The isolate was identified exclusively in intubated infants, occurring in five BPD^+^ Int^+^ infants and one BPD^-^ Int^+^ infant and was not detected in any BPD^-^ Int^-^ infants (Figure 6B). Nucleotide diversity analysis indicated that AU12-03-positive samples from BPD^+^ Int^+^ infants harbored genetically homogeneous populations, indicating infants likely harbored only the AU12-03 strain, whereas a single BPD^-^ Int^+^ infant exhibited higher diversity, consistent with the presence of multiple related *Staphylococcus epidermidis* strains (Figure 6A).

**Figure 6.**
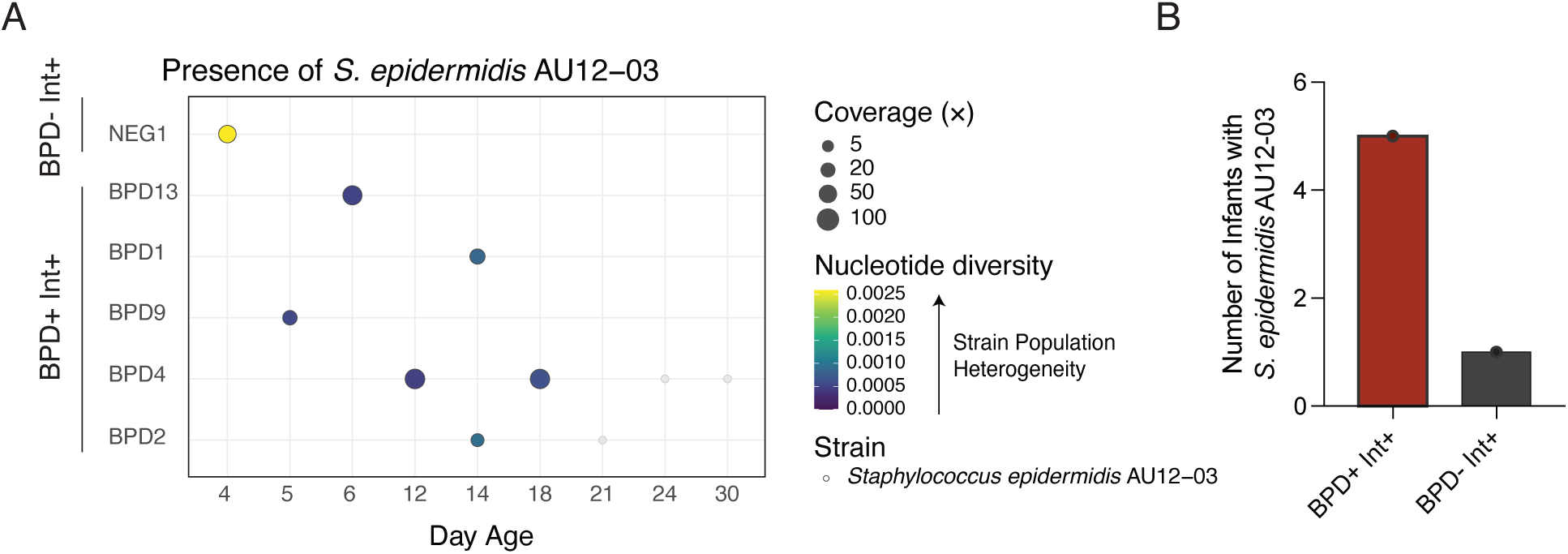
Detection of isolated strains from the NICU across infants. (A) Detection of *Staphylococcus epidermidis* AU12-03 in BPD^+^ Int^+^ and BPD^-^ Int^+^ infant stool metagenomes at various days of life filtered to 95% breadth 5X coverage, and 0.999 average nucleotide identity. (B) Instances of *Staphylococcus epidermidis* AU12-03 occurring in BPD^+^ Int^+^ or BPD^-^ Int^+^ infants.

### Temporal changes in the gut microbiome of individual infants

Changes in the gut microbiome over the first weeks of life varied between infants but often aligned with changes in feeding mode or antibiotic exposure. In two infants, early microbial communities were dominated by skin-associated microbes (e.g. *Staphylococcus* species) that shifted towards a composition dominated by enteric bacteria following the cessation of TPN and initiation of enteric feeding (Supplemental Figures 3 and 4). Antibiotic exposure also aligned with gut microbiome shifts in three infants, although variability between individuals and the sample size limit generalizability (Supplemental Figures 3 and 4).

## Discussion

This study provides new insights into the early development of the lung and gut microbiomes of premature infants, and how these communities relate to BPD and clinical exposures such as intubation, nutrition, and antibiotics. By using shotgun metagenomic sequencing of both tracheal aspirates and stool samples, we identified distinct microbial signatures associated with BPD and demonstrated that *Staphylococcus epidermidis* is a dominant and shared species in both sites among infants who developed BPD. These findings support the concept that early–life microbial colonization patterns may contribute to BPD pathogenesis and highlight the dynamic interplay between the lung and gut ecosystems during critical phases of neonatal development.

### Gut microbiome composition reflects clinical status and intervention

The gut microbiome composition varied significantly with both BPD status and intubation. Several individual taxa differed in relative abundance, suggesting specific microbial functions may contribute to shared phenotypes. Infants who went on to develop BPD and were intubated (BPD^+^ Int^+^) harbored a significantly higher abundance of skin-associated genera, including *Staphylococcus*, *Corynebacterium*, and *Cutibacterium*, compared to non-intubated, BPD-negative infants (BPD^-^ Int^-^), as well as intubated BPD-negative infants (BPD^-^ Int^+^). *Staphylococcus* species are pervasive in hospital environments – surveillance of *Staphylococcus aureus* strains have shown spatial and temporal proximity facilitate transmission between infants.^20^ These findings may also reflect cross-site seeding from the skin or from medical equipment more commonly utilized in intubated infants. The species-sharing events between the lung and gut within the same infant support a lung-gut axis of microbial exchange. Overall, these findings indicate that skin-associated flora, such as *Staphylococcus epidermidis*, may be associated with BPD development in the context of prior intubation.

Differential abundance analyses further highlighted microbial signatures associated with disease status. Several gut species, including *Enterococcus faecalis, Klebsiella aerogenes,* and *Klebsiella grimontii* showed differential abundance between groups, with *Klebsiella grimontii* enriched in BPD^-^ Int^-^ infants, *Klebsiella aerogenes* enriched in BPD^-^ Int^+^ infants, and *Enterococcus faecalis* enriched in both BPD^+^ Int^+^ and BPD^-^ Int^+^ infants. In contrast, *Staphylococcus epidermidis* was significantly enriched in the BPD^+^ Int^+^ group, supporting its potential role as a hallmark species of dysbiosis in this high-risk population.

### Overlap between lung and gut microbiomes in intubated infants

Microbial communities in the lung are naturally sparse therefore demanding high-resolution analysis. ^21,22^ The k-mer based alignment and reference-based alignment methodologies, as utilized here, are uniquely advantageous given strict parameters to identify truly “present” microbes within a low-biomass sample. Using these techniques among infants with sequencing data from both tracheal aspirate and stool samples, several species were detected in both lung and gut microbiomes, with *Staphylococcus epidermidis* being the most shared species. Additionally, comparative genomic analysis of *Staphylococcus epidermidis* isolates from infant TA and stool samples revealed strong evidence of species sharing. These events provide evidence of microbial overlap between the two mucosal sites. In intubated infants, mechanisms such as micro–aspiration, mucosal barrier immaturity, or shared environmental exposures may facilitate bidirectional microbial exchange. Despite this overlap, the lung and gut communities remained largely distinct, suggesting that shared species represent selective translocation or niche–specific expansion rather than wholesale community transfer. These findings support a potential microbiome–mediated lung–gut axis in the pathophysiology of BPD.

### Clinical interventions shape microbial dynamics

Longitudinal analysis revealed that both feeding practices and antibiotic administration were closely associated with shifts in gut and lung microbial composition. Transitions from total parenteral nutrition to enteral feeds, especially breastmilk, often coincided with changes in community structure, frequently involving an increase in opportunistic gut species. Similarly, antibiotic exposure led to notable decreases in species such as *Bacteroides thetaiotaomicron* and *Staphylococcus epidermidis*, while sometimes allowing expansion of other taxa like *Klebsiella pneumoniae* or *Enterococcus faecalis*. These findings underscore the sensitivity of the neonatal microbiome to external perturbations and support the hypothesis that early-life interventions can have profound effects on microbial ecology.

### *Staphylococcus epidermidis* is associated with bronchopulmonary dysplasia

The association of BPD progression and *Staphylococcus epidermidis* presence in both the lung and gut microbiomes of infants, suggests its potential role as a marker of dysbiosis and a contributor to disease pathogenesis. *Staphylococcus epidermidis—*a pervasive microbe commonly found on the skin, and in the respiratory and gastrointestinal tracts—is a major cause of late-onset neonatal sepsis.^23^ Neonates with *Staphylococcus epidermidis* sepsis are more likely to go on to develop bronchopulmonary dysplasia.^24,25^ The colonization of neonates with *Staphylococcus epidermidis* has been shown to occur through contact with caregivers, medical equipment, enteral feeding tubes,^26^and maternal breast milk.^23,27,28^ Emerging evidence suggests a strong link between *S. epidermidis* sepsis and inflammation-associated complications in preterm infants such as necrotizing enterocolitis, white matter injury, and bronchopulmonary dysplasia.^24,29^ Studies demonstrate that *S. epidermidis* activates an innate immune response by inducing the production of pro-inflammatory cytokines such as TNF-α, IL-1β, IL-6, and IL-8, ^30,31^ which, when combined with other antenatal and postnatal insults sustained by premature infants, may contribute to impaired lung development. Additionally, *Staphylococcus epidermidis* strains have been found to directly elicit pro-inflammatory signaling pathways in human lung epithelial cells,^32,33^ further suggesting a mechanistic link between increased abundance of *Staphylococcus epidermidis* and the development of bronchopulmonary dysplasia.

Limitations to this study includes its single center design, small sample size, and absence of a comparison group comprised of non-intubated BPD infants. Additionally, this study does not account for genetic propensities that may influence a neonate’s vulnerability to exogenous insults. The study is strengthened by the use of shotgun metagenomic sequencing that allows deeper analysis of the microbial communities. In contrast to prior studies that employed 16S rRNA gene sequencing, this approach enables species-level microbial identification, resulting in a more accurate representation of the true microbial composition. Future studies should assess the *Staphylococcus*-dependent mechanistic association between dysbiosis, immunologic maturation and response, and the development of bronchopulmonary dysplasia. Antibiotic-independent mechanisms of altering gut communities for shifting *Staphylococcus* dominance in susceptible lung and gut colonized preterm infants setting is also an important point of study in the development of potential preventative interventions. Moreover, longitudinal, multi-omic studies could elucidate causal relationships between microbial shifts and disease progression. Overall, this study demonstrates an association between lung and gut dysbiosis in premature infants and the development of bronchopulmonary dysplasia. *Staphylococcus epidermidis* emerged as a dominant species in both the lung and gut of infants who went on to develop BPD, potentially reflecting its prominence in the NICU environment^20^ and association with immune modulation. While our data cannot establish causality, the co-occurrence of specific taxa in the lung and gut, and their enrichment in BPD^+^ Int^+^ infants, raise the hypothesis that microbial composition and strain-level features may modulate airway injury and repair, thereby influencing BPD trajectories.

In conclusion in preterm infants, BPD and intubation status are associated with compositional shifts in the gut microbiome and with detectable, low–biomass lung communities enriched for select taxa, alongside within–infant species sharing between lung and gut. The results underscore a potential microbiome–mediated lung–gut axis in BPD and highlight clinical exposures, especially feeding and antibiotics, as important modulators of early colonization. These observations set the stage for mechanistic, exposure–aware, strain–resolved studies to determine whether modifying microbial communities can mitigate airway injury and improve outcomes in BPD.

## Acknowledgements

We thank the Genomics Facility of the Biotechnology Resource Center (BRC) of Cornell Institute of Biotechnology for their help with sequencing experiments. We would like to thank all participants in this research study and the Division of Neonatology at Weill Cornell Medicine for their continued support.

## Author Contributions

ZS performed sample collection, designed the data collection instruments, and collected data. SR and JJ performed sample collection. AH collected samples, and supervised data collection. ME performed experiments. ME and ST performed the data analysis. ZS, ME, ST, MZ, DK, SW, ELJ, and AH conceptualized and designed the study, and interpreted the data. All authors contributed to the drafting of the manuscript.

