## Supplemental Figures 1-4 and Tables 1-2 for "Increased *S. epidermidis* in the airway-gut microbiome of infants with bronchopulmonary dysplasia"

Supplemental Figures 1 – 4
Supplemental Tables 1 – 2

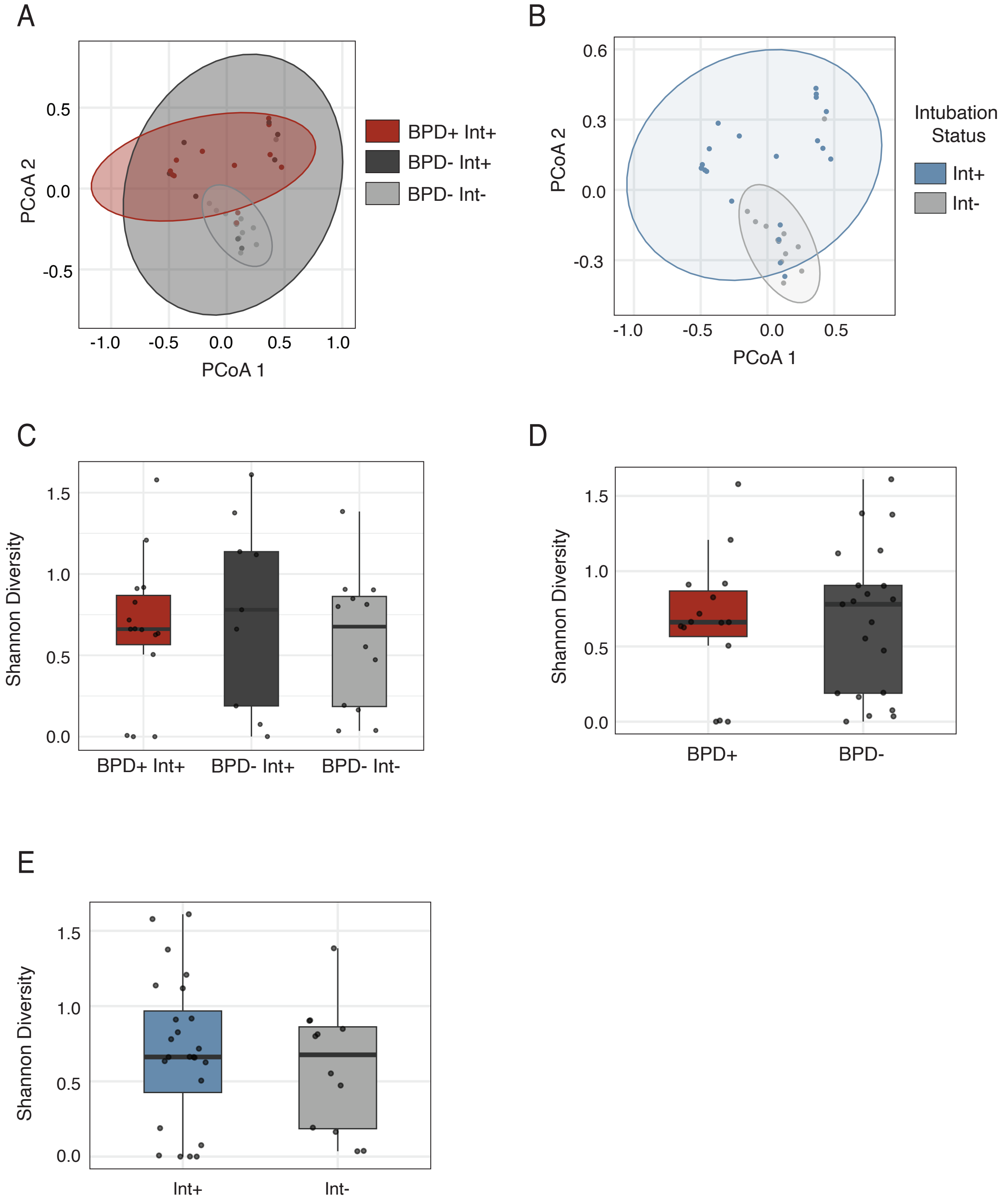


**Supplemental Figure 1 – Alpha and beta diversity analysis by BPD and intubation status.**

(A) Bray-Curtis dissimilarity among BPD$^+$ Int$^+$, BPD$^-$ Int$^+$, and BPD$^-$ Int$^-$ infants ($R^2 = 0.094$, F = 1.72, p = 0.022), (B) Int$^+$ and Int$^-$ infants (PERMANOVA: $R^2 = 0.072$, F. = 2.636, p = 0.007) using the first sample per infant. (C) Shannon diversity among BPD$^+$ Int$^+$, BPD$^-$ Int$^+$, and BPD$^-$ Int$^-$ infants (Kruskal-Wallis: $\chi^2 = 0.625$, df = 2, p = 0.73), (D) BPD$^+$ and BPD$^-$ infants (Wilcoxon rank-sum: W = 148, p = 0.773), and (E) Int$^+$ and Int$^-$ infants (Wilcoxon rank-sum: W = 156, p = 0.6995) using the first sample per infant.

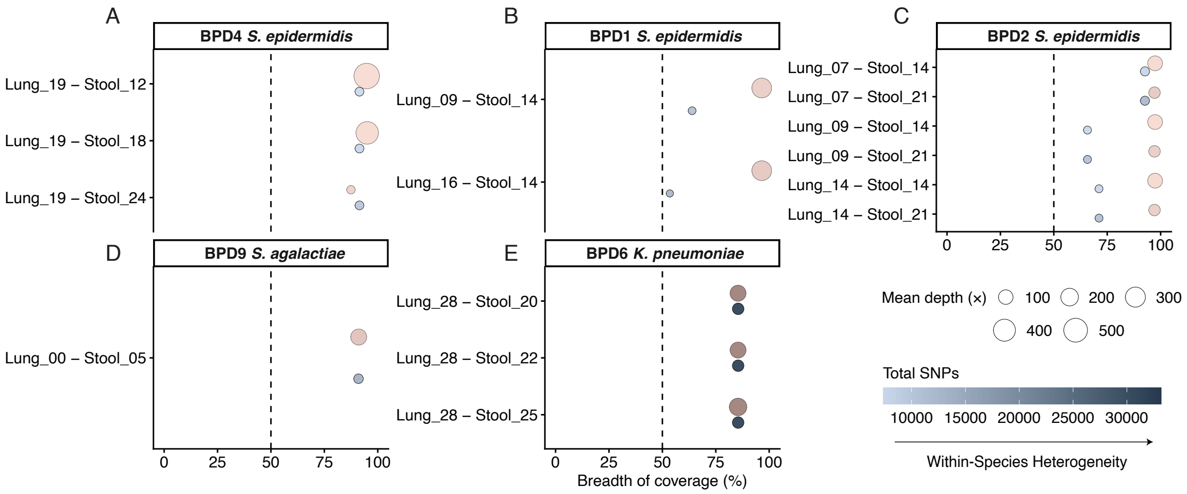


**Supplemental Figure 2 – Species sharing analysis using minimap2 between lung and stool metagenomes.** (A) Infant – BPD4, (B) Infant – BPD1, (C) Infant – BPD2, (D) Infant – BPD9, and (E) Infant – BPD6 at different days of life. On the x-axis is the breadth of coverage between the lung and stool metagenomes and the given species reference genome. Also represented on the x-axis is a dotted line denoting a threshold of 50% breadth of coverage. Color of the point represents the site (red = stool, blue = lung), and size of the point represents the depth of coverage of the alignment between lung, stool, and the given species genome.

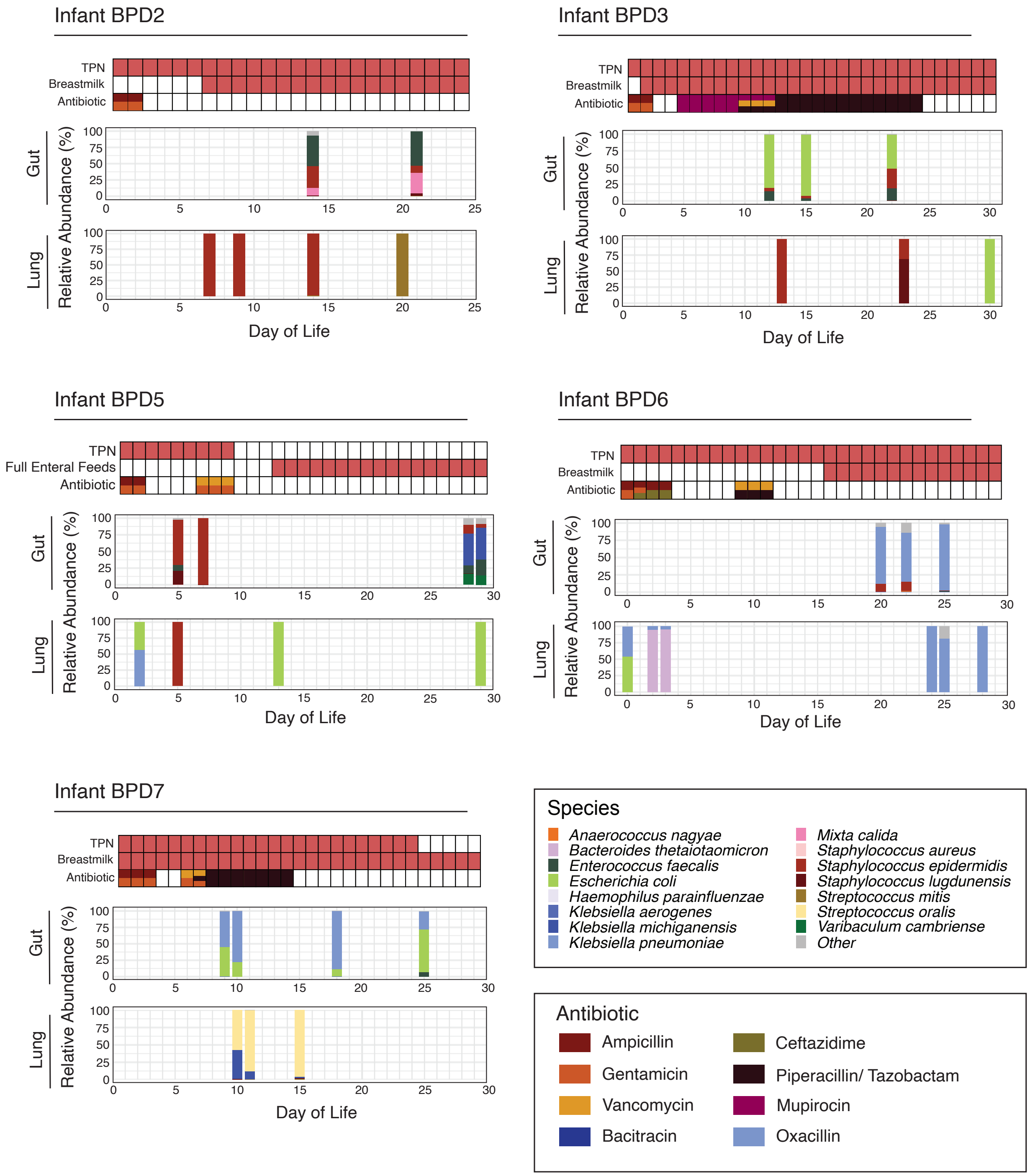


**Supplemental Figure 3 – Microbial community shifts in the gut and lower airway microbiome of individual infants over periods of feeding changes and antibiotic intake.** The top 15 species within the samples plotted are denoted in the legend with the other species combined into the "Other" category. If two samples were collected for one infant on the same day, the first sample was plotted.

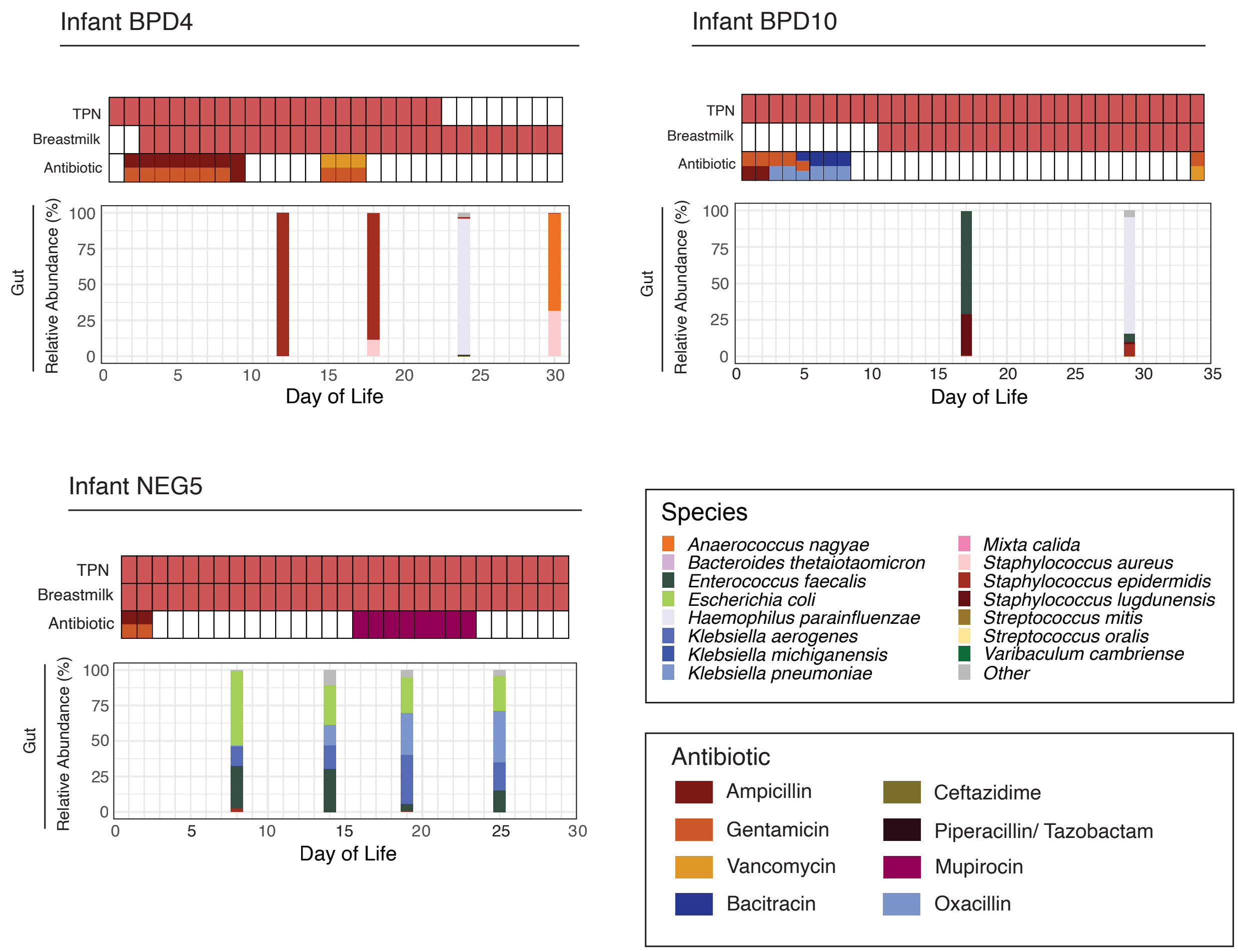


**Supplemental Figure 4– Microbial community shifts in the gut microbiome of individual infants over periods of feeding changes and antibiotic intake.** The top 15 species within the samples plotted are denoted in the legend with the other species combined into the “Other” category. If two samples were collected for one infant on the same day, the first sample was plotted.

**Supplemental Table 1 –** Species tracking alignments and their NCBI RefSeq IDs.

| query1 | query2 | lineage | NCBI RefSeq |
|---|---|---|---|
| lung_BPD9_0 | stool_BPD9_5 | *Streptococcus agalactiae* | GCF_001552035.1 |
| lung_BPD2_7 | stool_BPD2_21 | *Staphylococcus epidermidis* | GCF_006094375.1 |
| lung_BPD2_9 | stool_BPD2_21 | *Staphylococcus epidermidis* | GCF_006094375.1 |
| lung_BPD6_0 | stool_BPD6_22 | *Klebsiella pneumoniae* | GCF_000240185.1 |
| lung_BPD6_28 | stool_BPD6_22 | *Klebsiella pneumoniae* | GCF_000240185.1 |
| lung_BPD6_28 | stool_BPD6_25 | *Klebsiella pneumoniae* | GCF_000240185.1 |
| lung_BPD6_28 | stool_BPD6_25_timepoint_2 | *Klebsiella pneumoniae* | GCF_000240185.1 |
| lung_BPD2_9 | stool_BPD2_14 | *Staphylococcus epidermidis* | GCF_006094375.1 |
| lung_BPD2_7 | stool_BPD2_14 | *Staphylococcus epidermidis* | GCF_006094375.1 |
| lung_BPD2_14 | stool_BPD2_21 | *Staphylococcus epidermidis* | GCF_006094375.1 |
| lung_BPD6_0 | stool_BPD6_25_timepoint_2 | *Klebsiella pneumoniae* | GCF_000240185.1 |
| lung_BPD6_0 | stool_BPD6_25 | *Klebsiella pneumoniae* | GCF_000240185.1 |
| lung_BPD4_19 | stool_BPD4_12 | *Staphylococcus epidermidis* | GCF_006094375.1 |
| lung_BPD5_5 | stool_BPD5_28 | *Staphylococcus epidermidis* | GCF_006094375.1 |
| lung_BPD4_19 | stool_BPD4_24 | *Staphylococcus epidermidis* | GCF_006094375.1 |
| lung_BPD5_5 | stool_BPD5_29 | *Staphylococcus epidermidis* | GCF_006094375.1 |
| lung_BPD2_14 | stool_BPD2_14 | *Staphylococcus epidermidis* | GCF_006094375.1 |
| lung_BPD6_28_timepoint_2 | stool_BPD6_22 | *Klebsiella pneumoniae* | GCF_000240185.1 |
| lung_BPD3_13 | stool_BPD3_15 | *Staphylococcus epidermidis* | GCF_006094375.1 |
| lung_BPD6_28_timepoint_2 | stool_BPD6_25 | *Klebsiella pneumoniae* | GCF_000240185.1 |
| lung_BPD6_28 | stool_BPD6_20 | *Klebsiella pneumoniae* | GCF_000240185.1 |
| lung_BPD3_30 | stool_BPD3_15 | *Escherichia coli* | GCF_000005845.2 |
| lung_BPD6_28_timepoint_2 | stool_BPD6_25_timepoint_2 | *Klebsiella pneumoniae* | GCF_000240185.1 |
| lung_BPD3_30 | stool_BPD3_15_timepoint_2 | *Escherichia coli* | GCF_000005845.2 |
| lung_BPD6_2 | stool_BPD6_25 | *Klebsiella pneumoniae* | GCF_000240185.1 |
| lung_BPD6_2 | stool_BPD6_25_timepoint_2 | *Klebsiella pneumoniae* | GCF_000240185.1 |
| lung_BPD5_5 | stool_BPD5_7 | *Staphylococcus epidermidis* | GCF_006094375.1 |
| lung_BPD6_28_timepoint_2 | stool_BPD6_20 | *Klebsiella pneumoniae* | GCF_000240185.1 |
| lung_BPD4_19 | stool_BPD4_18 | *Staphylococcus epidermidis* | GCF_006094375.1 |
| lung_BPD5_29 | stool_BPD5_29 | *Escherichia coli* | GCF_000005845.2 |
| lung_BPD6_0 | stool_BPD6_20 | *Klebsiella pneumoniae* | GCF_000240185.1 |
| lung_BPD6_2 | stool_BPD6_22 | *Klebsiella pneumoniae* | GCF_000240185.1 |
| lung_BPD3_13 | stool_BPD3_15_timepoint_2 | *Staphylococcus epidermidis* | GCF_006094375.1 |
| lung_BPD6_3 | stool_BPD6_25 | *Klebsiella pneumoniae* | GCF_000240185.1 |
| lung_BPD6_3 | stool_BPD6_25_timepoint_2 | *Klebsiella pneumoniae* | GCF_000240185.1 |
| lung_BPD6_24 | stool_BPD6_25 | *Klebsiella pneumoniae* | GCF_000240185.1 |
| lung_BPD3_23 | stool_BPD3_15 | *Staphylococcus epidermidis* | GCF_006094375.1 |
| lung_BPD4_19_timepoint_2 | stool_BPD4_18 | *Staphylococcus aureus* | GCF_000013425.1 |

| | | | |
|---|---|---|---|
| lung_BPD6_3 | stool_BPD6_22 | *Klebsiella pneumoniae* | GCF_000240185.1 |
| lung_BPD3_30 | stool_BPD3_22 | *Escherichia coli* | GCF_000005845.2 |
| lung_BPD6_24 | stool_BPD6_22 | *Klebsiella pneumoniae* | GCF_000240185.1 |
| lung_BPD6_24 | stool_BPD6_25_timepoint_2 | *Klebsiella pneumoniae* | GCF_000240185.1 |
| lung_BPD6_2 | stool_BPD6_20 | *Klebsiella pneumoniae* | GCF_000240185.1 |
| lung_BPD7_11 | stool_BPD7_25 | *Klebsiella michiganensis* | GCF_015139575.1 |
| lung_BPD6_25_timepoint_2 | stool_BPD6_25 | *Klebsiella pneumoniae* | GCF_000240185.1 |
| lung_BPD3_30 | stool_BPD3_12 | *Escherichia coli* | GCF_000005845.2 |
| lung_BPD5_29 | stool_BPD5_28 | *Escherichia coli* | GCF_000005845.2 |
| lung_BPD7_10 | stool_BPD7_25 | *Klebsiella michiganensis* | GCF_015139575.1 |
| lung_BPD3_23 | stool_BPD3_15_timepoint_2 | *Staphylococcus epidermidis* | GCF_006094375.1 |
| lung_BPD6_25_timepoint_2 | stool_BPD6_22 | *Klebsiella pneumoniae* | GCF_000240185.1 |
| lung_BPD7_1 | stool_BPD7_25 | *Klebsiella pneumoniae* | GCF_000240185.1 |
| lung_BPD6_25_timepoint_2 | stool_BPD6_25_timepoint_2 | *Klebsiella pneumoniae* | GCF_000240185.1 |
| lung_BPD1_9 | stool_BPD1_14 | *Staphylococcus epidermidis* | GCF_006094375.1 |
| lung_BPD6_3 | stool_BPD6_20 | *Klebsiella pneumoniae* | GCF_000240185.1 |
| lung_BPD7_4 | stool_BPD7_25 | *Klebsiella pneumoniae* | GCF_000240185.1 |
| lung_BPD4_19_timepoint_2 | stool_BPD4_30 | *Staphylococcus aureus* | GCF_000013425.1 |
| lung_BPD6_25 | stool_BPD6_25 | *Klebsiella pneumoniae* | GCF_000240185.1 |
| lung_BPD6_25 | stool_BPD6_22 | *Klebsiella pneumoniae* | GCF_000240185.1 |
| lung_BPD6_25_timepoint_2 | stool_BPD6_20 | *Klebsiella pneumoniae* | GCF_000240185.1 |
| lung_BPD1_16 | stool_BPD1_14 | *Staphylococcus epidermidis* | GCF_006094375.1 |

**Supplemental Table 2 –** *Staphyloccocus epidermidis* strains utilized in strain tracking analysis.

| Strain | NCBI RefSeq |
|---|---|
| *Staphyloccocus epidermidis* ATCC 14990 | GCF_006094375.1 |
| *Staphyloccocus epidermidis* B1276912 | GCF_019329745.1 |
| *Staphyloccocus epidermidis* B1272014 | GCF_019329665.1 |
| *Staphyloccocus epidermidis* UMCG345 | GCF_045940975.1 |
| *Staphyloccocus epidermidis* UMCG350 | GCF_045939605.1 |
| *Staphyloccocus epidermidis* B1230143 | GCF_019329425.1 |
| *Staphyloccocus epidermidis* AR4072 | GCF_031326165.1 |
| *Staphyloccocus epidermidis* IRL01 | GCF_009685135.1 |
| *Staphyloccocus epidermidis* B1266875 | GCF_019329585.1 |
| *Staphyloccocus epidermidis* TMDU-265 | GCF_024204785.1 |
| *Staphylococcus hominis* FDAARGOS_575 | GCF_003812505.1 |
| *Staphylococcus capitis* DSM 6717 | GCF_040739365.1 |
| *Staphylococcus epidermidis* AU12-03 | NA |
